# Engineered CRISPR-Base Editors as a Permanent Treatment for Familial Dysautonomia

**DOI:** 10.1101/2024.11.27.625322

**Authors:** Shuqi Yun, Anil Chekuri, Jennifer Art, Krishnakanth Kondabolu, Samikshya Pokharel, Paolo Pigini, Sydnee Dymock, David Rufino-Ramos, Susan A. Slaugenhaupt, Nadja Zeltner, Benjamin P. Kleinstiver, Elisabetta Morini, Christiano R. R. Alves

## Abstract

Familial dysautonomia (FD) is a fatal autosomal recessive sensory and autonomic neuropathy. FD is caused by a T-to-C point mutation in intron 20 of the *Elongator acetyltransferase complex subunit 1* (*ELP1*) gene, which results in tissue-specific skipping of exon 20 to cause a premature termination codon and thus redues ELP1 protein levels. Here, we developed a CRISPR-Cas-based cytosine base editing strategy to permanently correct the disease-causing mutation and restore canonical mRNA splicing. Through systematic engineering of base editors and guide RNAs, we identified an optimal editor configuration capable of achieving up to 70% on-target correction in human cells and that restored *ELP1* exon 20 inclusion. To enable in vivo delivery, a dual adeno-associated virus (AAV) intein-split base editor was delivered via intravenous injections in a humanized FD mouse mode, resulting in genetic correction and significantly increased *ELP1* exon 20 inclusion in the brain and other tissues. In FD patient-derived iPSC-sympathetic neurons, we observed ∼10% correction efficiency but rescued disease-associated neuronal hyperactivity, demonstrating that partial correction can restore functional phenotypes. Genome-wide analyses revealed minimal off-target editing across multiple human cell types, supporting the specificity of this approach. Together, these findings establish a precise and permanent genome editing strategy for FD and supports the development of a one-time disease-modifying therapy for FD and highlights the therapeutic potential of base editing for splicing disorders.

## Introduction

Familial dysautonomia (FD), also known as hereditary sensory and autonomic neuropathy type III (HSANIII) or Riley-Day syndrome, is a rare autosomal recessive congenital neurodevelopmental disorder characterized by reduced developmental survival and progressive degeneration of sensory and autonomic neurons. FD is caused by a T-to-C transition at the 6th position of intron 20 (c.2204+6T>C) in the Elongator acetyltransferase complex subunit 1 (*ELP1*, previously known as *IKBKAP*) (*1, 2*). This mutation (hereafter referred to as *ELP1* T6C) results in tissue-specific skipping of exon 20, leading to a premature termination codon and reduced ELP1 protein levels (*1, 3*). The most pronounced reduction of ELP1 protein occurs in the central and peripheral nervous systems (*3*), and >99.5% of FD patients are homozygous, with no known disease manifestations in heterozygous carriers (*1, 4*). Therefore, therapeutic strategies aimed at correcting this specific splicing defect have the potential to provide clinical benefit across the entire FD patient population.

Clinically, FD is characterized by impaired pain and temperature sensation, orthostatic hypotension, tachycardia, labile blood pressure, gait ataxia, and progressive optic neuropathy (*5–10*) (**Fig. 1A**). Recurrent aspiration due to neurogenic dysphagia contributes to chronic pulmonary disease . Patients also experience brisk episodes of severe hypertension, tachycardia, skin blotching, retching, and vomiting, referred to as autonomic crises. Unexplained sudden death, aspiration pneumonia, and respiratory insufficiency remain the leading causes of death (*5, 11*). Currently, there are no FDA-approved disease-modifying therapies capable of slowing or reversing the progressive neuronal degeneration in FD.

**Fig. 1.**
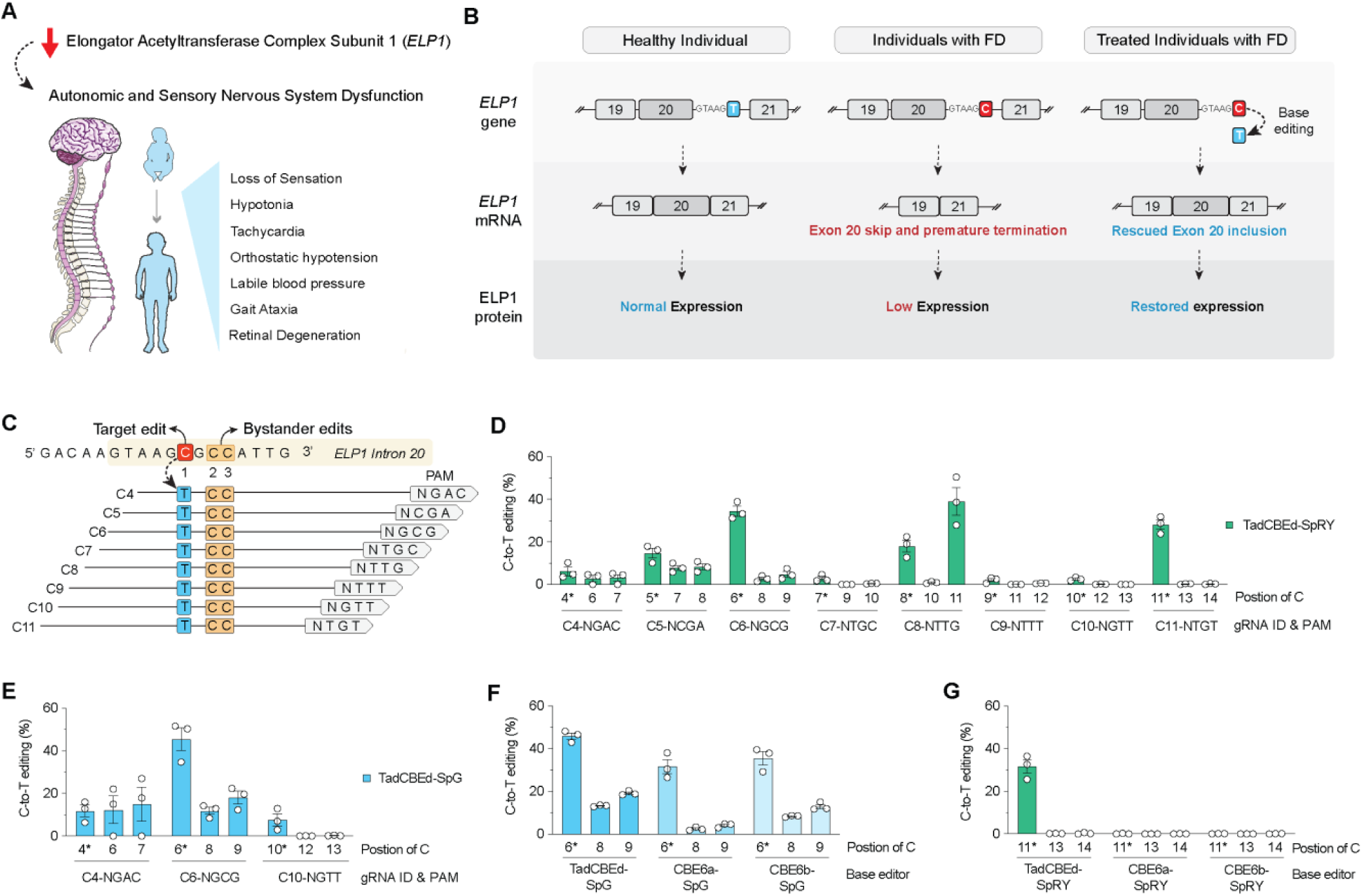
Engineering cytosine base editors (CBEs) to correct the pathogenic ELP1 T6C mutation associated with familial dysautonomia. **(A)** Schematic illustrating major clinical manifestations of familial dysautonomia (FD) caused by reduced ELP1 expression and dysfunction of the autonomic and sensory nervous systems. **(B)** Schematic of the FD-associated ELP1 intronic T-to-C mutation (T6C) at position +6 of intron 20, which promotes exon 20 skipping, reduces ELP1 protein expression, and is targeted for correction by cytosine base editing. **(C)** Schematic of the ELP1 target region showing guide RNA (gRNA) target sites and PAM sequences used for cytosine base editor screening. The pathogenic cytosine is highlighted in red and adjacent potential bystander editing sites are highlighted in orange. (**D** and **E**) On-target C-to-T editing efficiencies across the ELP1 locus in HEK293T-ELP1-TC6 cells using TadCBEd fused to PAM-flexible SpCas9 variants SpRY (**D**) or SpG (**E**) paired with distinct gRNAs. Editing frequencies were quantified by targeted deep sequencing and analyzed using CRISPResso2. (**F** and **G**) Comparison of editing efficiencies obtained using TadCBEd, CBE6a, or CBE6b deaminase domains fused to SpG paired with gRNA-C6 (**F**) or SpRY paired with gRNA-C11 (**G**). Data in panels **D** to **G** represent mean ± SEM with individual data points shown; n = 3 independent biological replicates.

Previous efforts to develop therapeutic approaches for FD include splicing modulator compounds, antisense oligonucleotides (ASOs), modified exon-specific U1 small nuclear RNAs, and gene replacement therapy (*12–18*). However, none of these strategies provide a permanent cure or have been FDA-approved. ASOs, modified exon-specific U1 small nuclear RNAs, and gene replacement therapies are constrained by transient effects, along with uncertain longevity of expression, utilization of ubiquitous exogenous promoters, and delivery constraints. Splicing modulator compounds, while preferred due to the ease of oral administration, will require lifelong treatment. Development of a genome editing technology capable of permanently correcting the underlying FD splicing mutation to adequately maintain endogenous expression of *ELP1* would overcome many of these limitations.

Base editors (BEs) and prime editors (PEs) are genome editing technologies capable of introducing point mutations (*19*). Recently, we reported pre-clinical efforts to optimize novel BE technologies to treat spinal muscular atrophy (SMA), a severe pediatric neuromuscular disease (*20*), and multisystemic smooth muscle dysfunction syndrome (MSMDS), a genetic vasculopathy associated with stroke and death in childhood (*21*). In a similar context, other recent pre-clinical studies have demonstrated the potential of BEs and PEs to treat neurological conditions (*22–25*). Together, this evidence highlights the strong therapeutic potential of customized BEs to safely and effectively introduce corrective genetic edits.

In this study, we investigated the potential of developing a genetic treatment for FD by correcting the *ELP1* TC6 mutation using cytosine base editing (CBE) (**Fig. 1B**). We systematically optimized a series of base editors and conducted a comprehensive evaluation of on-target and off-target editing profiles. These analyses revealed minimal off-target editing, demonstrating high specificity of these optimized base editors. To assess the *in vivo* efficacy of the optimized base editor strategy, we utilized a humanized FD mouse model harboring the human *ELP1* gene carrying the T-to-C mutation, as well as FD patient-derived induced pluripotent stem cell (iPSC)-differentiated sympathetic neurons. Our findings demonstrate that engineering base editors can effectively correct the *ELP1* splicing defects *in vivo* and in human neurons, leading to phenotypic rescue. This work provides the essential proof-of-concept data for advancing a durable, single-dose therapeutic strategy for FD.

## Results

### Generation of HEK 293T cell lines harboring the major FD-associated *ELP1* T6C mutation

We first established a homozygous HEK293T cell line harboring the *ELP1* T6C mutation using adenosine base editing (ABE). This cell line was generated to enable rapid screening and prioritization by correction efficiency of editor and guide RNA (gRNA) combinations. We initially screened multiple combinations of ABEs and gRNAs targeting the *ELP1* locus and identified several conditions achieving >50% precise introduction of the pathogenic mutation in HEK293T cells, with variable levels of bystander editing (**Fig. S1A**). Based on these results, we selected ABE8.20-SpRY paired with gRNA A6 as the most efficient and precise base editing strategy for generating a clonal screening platform. HEK293T cells were transfected with this editor configuration, followed by serial dilution and single-cell isolation (**Fig. S1B**). Targeted sequencing of the *ELP1* locus confirmed the generation of clones harboring homozygous *ELP1 T6C* mutations. To maximize sensitivity for subsequent correction studies, we selected a homozygous clone carrying the *ELP1 T6C* mutation without detectable bystander edits (hereafter referred to as HEK293T-*ELP1-TC6*). This engineered cell line was subsequently used throughout the study to systematically screen and optimize base editing strategies for correction of the FD mutation throughout the study.

### Development of cytosine base editors to correct the *ELP1* T6C mutation

Using the newly generated HEK293T-*ELP1-T6C* homozygous cell line, we investigated whether cytosine base editors (CBEs), which catalyze C•G-to-T•A conversions (*26*), could correct the pathogenic *ELP1 T6C* mutation. This target site presents several technical challenges for base editing. First, CBEs target cytosines within a narrow editing window positioned at a fixed distance from a protospacer adjacent motif (PAM), and the sequence adjacent to the *ELP1* mutation lacks canonical NGG PAMs recognized by wild-type (WT) SpCas9 and is therefore inaccessible to this enzyme (**Fig. 1C**). Second, the target cytosine is flanked by two adjacent cytosines, increasing the risk of bystander editing (**Fig. 1C**). To overcome these constraints, we leveraged engineered SpCas9 variants with expanded PAM compatibility, including SpRY that is near-PAMless motifs, and SpG that preferentially target sites with NGN PAMs (*27*). We additionally designed eight gRNAs spanning the *ELP1* target region to position the pathogenic cytosine at editing window positions C4–C11 (**Fig. 1C**).

We then evaluated SpRY or SpG nickases fused with different deaminases and found that CBEs encoding the recently developed TadCBEd deaminase (*26*) consistently outperformed conventional editors including BE4max (*28*) and prime editing, for correction of the major FD mutation (**Fig. 1D-E** and **Fig. S2A-C**). Based on these results, we selected TadCBEd-SpRY and TadCBEd-SpG as editor variants for further experiments.

TadCBEd-SpRY mediated efficient C-to-T editing across nearly all tested gRNAs within the editable window, with the highest precision observed using gRNA-C11 (NTG PAM), which achieved efficient correction without detectable bystander editing (**Fig. 1D**). In parallel, TadCBEd-SpG also enabled robust editing across all compatible NGN PAM-containing gRNAs (**Fig. 1E**). Among all conditions tested, TadCBEd-SpG paired with gRNA-C6 (NGC PAM) produced the highest (∼50%) overall editing efficiency (**Fig. 1E**). Although this configuration generated bystander edits at two adjacent cytosines, we further demonstrate in this manuscript that these edits were confined to the intronic region and therefore did not alter the *ELP1* coding sequence or negatively effect *ELP1* exon 20 inclusion. Furthermore, the TadCBEd deaminase demonstrated superior activity compared to other highly active deaminases, such as CBE6a and CBE6b (*29*), across multiple configurations (**Fig. 1F-G**). Collectively, these findings demonstrate that engineered CBEs can efficiently and precisely correct the ELP1 T6C mutation in human cells.

### Optimization of an AAV-CBE strategy to correct the *ELP1* T6C mutation

Based on our initial screening studies, we prioritized two editor configurations for further development: (1) TadCBEd-SpRY paired with gRNA-C11, which enabled precise on-target editing without detectable bystander edits, and (2) TadCBEd-SpG paired with gRNA-C6, which produced the highest overall editing efficiency. To facilitate *in vivo* delivery, we engineered and produced adeno-associated virus (AAV)-compatible intein-split base editor systems capable of packaging these large constructs (*20*).

We first generated intein-split architectures containing gRNA expression cassettes positioned either exclusively in the C terminal vector or distributed across both N- and C-terminal vectors (**Fig. 2A**). Both configurations promoted efficient on-target editing activity in HEK293T-*ELP1-TC6* cells (**Fig. 2B-C**). Notably, delivery of the TadCBEd-SpG/gRNA-C6 configuration through the intein-split system resulted in a substantial increase in *ELP1* exon 20 inclusion, whereas the TadCBEd-SpRY/gRNA-C11 combination did not produce splicing rescue despite detectable on-target editing (**Fig. 2D-F**). These findings suggest that maximizing editing efficiency may be more important for functional splicing correction than minimizing bystander editing at this locus. Moreover, the adjacent intronic bystander edits generated by the TadCBEd-SpG/gRNA-C6 configuration may further contribute to improved exon 20 inclusion, potentially through modulation of local splicing regulatory elements.

**Fig. 2.**
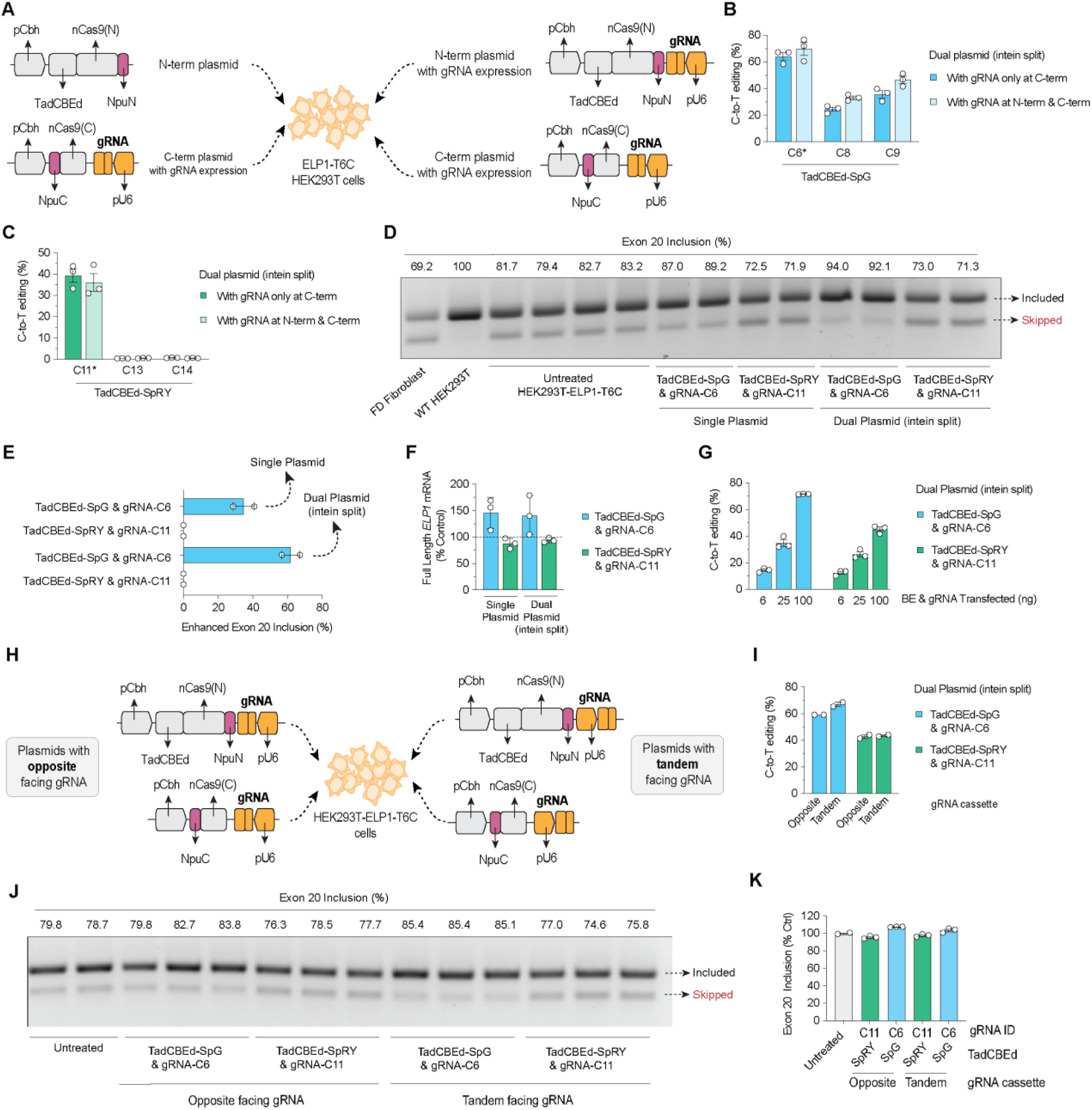
Engineering and optimization of intein-split AAV-compatible base editor systems for correction of the ELP1 T6C mutation. (**A**) Schematic of intein-split AAV plasmid architectures used for delivery of TadCBEd fused to SpG or SpRY in HEK293T-ELP1-TC6 cells. Constructs contained guide RNA (gRNA) expression cassettes positioned either exclusively within the C-terminal vector or in both N-terminal and C-terminal vectors. (**B** and **C**) On-target C-to-T editing efficiencies in HEK293T-ELP1-TC6 cells following transfection with intein-split TadCBEd-SpG (**B**) or TadCBEd-SpRY (**C**) AAV plasmid systems illustrated in (**A**). Editing frequencies were quantified by targeted deep sequencing and analyzed using CRISPResso2. **(D)** RT-PCR analysis of ELP1 exon 20 inclusion in HEK293T-ELP1-TC6 cells treated with single-plasmid or intein-split dual-plasmid base editor systems. Included and skipped ELP1 splice isoforms are indicated. **(E)** Quantification of exon 20 inclusion from experiments shown in (**D**). **(F)** RT-qPCR analysis of full-length ELP1 transcript expression following treatment with single-plasmid or intein-split dual-plasmid base editor systems containing TadCBEd-SpG paired with gRNA-C6 or TadCBEd-SpRY paired with gRNA-C11. **(G)** On-target C-to-T editing efficiencies following plasmid titration experiments using intein-split TadCBEd base editor systems in HEK293T-ELP1-TC6 cells. **(H)** Schematic comparing opposite-facing and tandem-facing guide RNA cassette orientations within intein-split AAV plasmid architectures. **(I)** Comparison of on-target editing efficiencies between opposite-facing and tandem-facing gRNA cassette configurations in HEK293T-ELP1-TC6 cells. **(J)** Representative RT-PCR analysis of ELP1 exon 20 inclusion following treatment with intein-split constructs shown in (H). **(K)** Quantification of exon 20 inclusion from experiments shown in (J). Data in panels **B**, **C**, **E** to **G**, **I**, and **K** represent mean ± SEM with individual data points shown; n = 2 or 3 independent biological replicates.

Because transgene expression from AAV vectors *in vivo* is often lower than that achieved with standard plasmid transfection, we next performed plasmid titration experiments to model reduced editor expression levels under more physiologically relevant conditions. Across all tested doses, TadCBEd-SpG paired with gRNA-C6 consistently achieved the highest levels of *ELP1* T6C correction (**Fig. 2G**).

We further investigated whether the orientation of the gRNA cassette relative to the split editor architecture influenced editing performance (**Fig. 2H**). Comparison of tandem-facing versus opposite-facing gRNA configurations revealed that tandem orientation modestly improved on-target editing efficiency. Importantly, both configurations resulted in restoration of *ELP1* exon 20 inclusion when using TadCBEd-SpG/gRNA-C6, but not with TadCBEd-SpRY/gRNA-C11 (**Fig. 2I-K**), consistent with our earlier observations. Taken together, these results show that we have established an AAV-compatible intein-split base editor system capable of efficient correction of the *ELP1 T6C* mutation and robust rescue of ELP1 splicing.

### Assessment of CBE-mediated *ELP1* T6C specificity

Because base editors can induce unintended genome-wide off-target edits, we next performed a comprehensive specificity assessment using both an unbiased cell-based detection assay (GUIDE-seq2) and in silico prediction analysis with Cas-OFFinder seq (*30–32*). For GUIDE-seq2, we used SpG nuclease paired with gRNA-C6 and SpRY nuclease paired with gRNA-C11 in HEK293T-*ELP1-T6C* cells.

Sequencing analysis confirmed robust on-target editing activity (**Fig. 3A**) and successful integration of the GUIDE-seq dsODN tag at cleavage sites (**Fig. 3B**). GUIDE-seq2 identified five and six candidate off-target sites for SpG/gRNA-C6 and SpRY/gRNA-C11, respectively (**Fig. 3C** and **Fig. S3A-B**). Notably, approximately 70% of GUIDE-seq2 reads generated with SpG/gRNA-C6 mapped to the intended *ELP1* target site, compared to ∼55% for SpRY/gRNA-C11 (**Fig. 3D**), suggesting greater targeting specificity and on-target preference for the SpG/gRNA-C6 configuration.

**Fig. 3.**
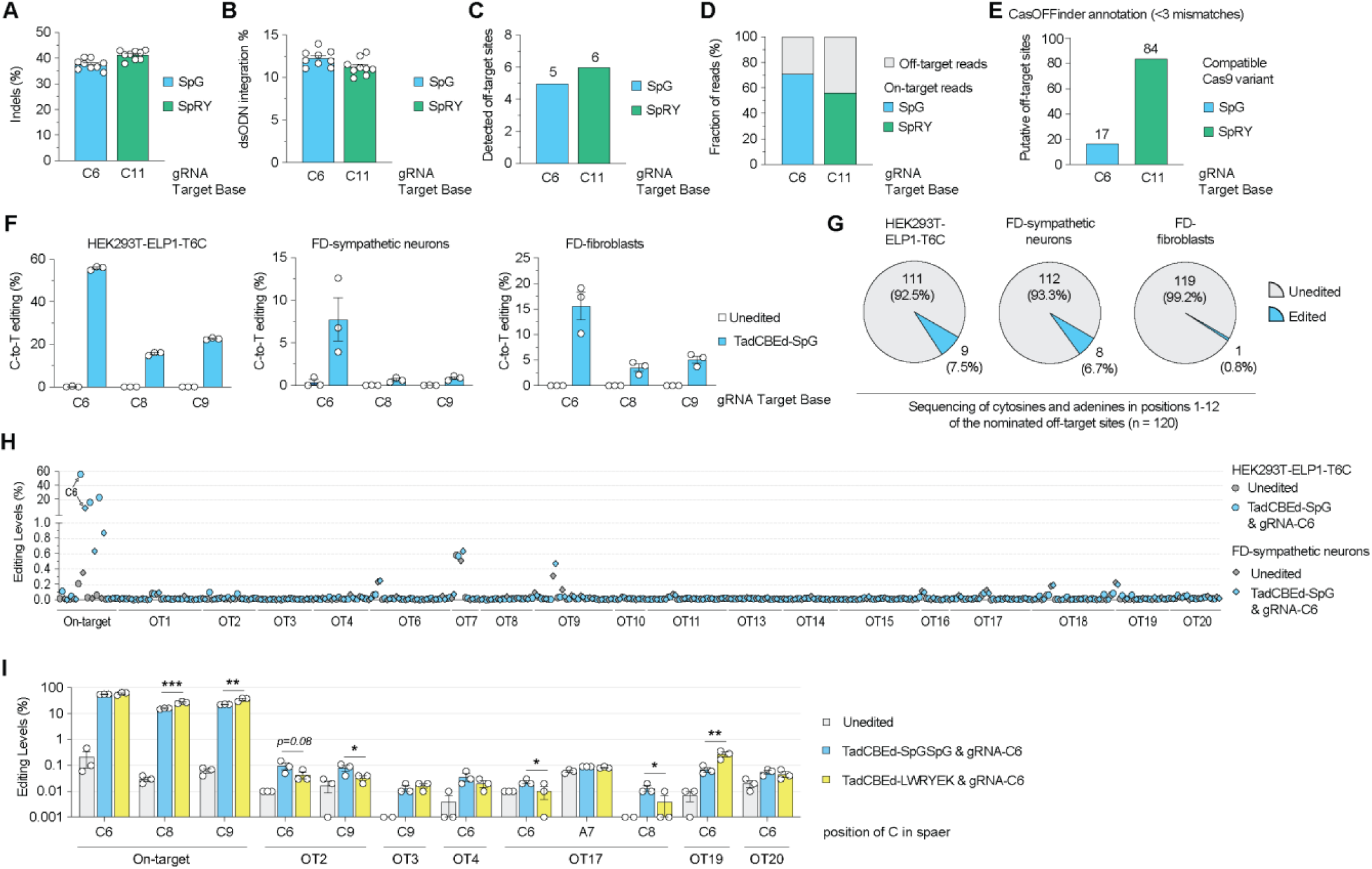
Specificity assessment of ELP1 base editing using GUIDE-seq2 and targeted off-target validation. (**A** and **B**) GUIDE-seq2 analysis in HEK293T-ELP1-TC6 cells using SpG nuclease paired with gRNA-C6 or SpRY nuclease paired with gRNA-C11. (**A**) Indel frequencies at the on-target ELP1 locus. (**B**) Quantification of reads containing integrated GUIDE-seq dsODN tags. **(C)** Total number of GUIDE-seq2-nominated off-target sites identified for SpG/gRNA-C6 and SpRY/gRNA-C11. **(D)** Percentage of GUIDE-seq2 reads mapping to the intended ELP1 target site or cumulative off-target loci for each nuclease/gRNA configuration. **(E)** Number of computationally predicted off-target sites containing up to three mismatches relative to gRNA-C6 or gRNA-C11 spacer sequences, identified using Cas-OFFinder. **(F)** On-target C-to-T editing efficiencies in HEK293T-ELP1-TC6 cells, FD patient-derived iPSC sympathetic neurons, and FD patient fibroblasts following treatment with TadCBEd-SpG paired with gRNA-C6. **(G)** Summary of validated off-target editing events detected across the three human cell types shown in (**F**). Pie charts indicate the number and percentage of loci without detectable editing among all nominated off-target sites identified by GUIDE-seq2 and Cas-OFFinder analyses. **(H)** Quantification of on-target and off-target base editing frequencies in HEK293T-ELP1-TC6 cells, FD patient-derived iPSC sympathetic neurons, and FD patient fibroblasts treated with TadCBEd-SpG and gRNA-C6. Genomic DNA from untreated and treated cells was analyzed by targeted deep sequencing of the on-target ELP1 locus and 19 nominated off-target sites using CRISPResso2. **(I)** Comparison of on-target and selected off-target editing frequencies using TadCBEd-SpG or the engineered PAM-selective TadCBEd-LWRYEK variant paired with gRNA-C6. Editing frequencies were quantified by targeted deep sequencing. Data in (**A**), (**B**), (**F**), (**H**), and (**I**) represent mean ± SEM with individual data points shown; n = 3 independent biological replicates unless otherwise indicated.

To complement these unbiased analyses, we next performed computational prediction of potential off-target sites containing up to three mismatches using Cas-OFFinder. This analysis identified 17 candidate sites for SpG/gRNA-C6 and 84 candidate sites for SpRY/gRNA-C11 (**Fig. 3E** and **Fig. S4A-C**). Two of the predicted SpG/gRNA-C6 off-target sites overlapped with GUIDE-seq2-identified sites, further supporting the specificity of this strategy.

Given the superior editing efficiency and targeting profile of the SpG/gRNA-C6 configuration, we prioritized this editor for downstream validation studies. We performed targeted deep sequencing across a combined set of 20 nominated off-target loci derived from GUIDE-seq2 and Cas-OFFinder analyses (**Table S1**). Validation steps were conducted in three distinct human cell types that demonstrated efficient on-target editing: HEK293T-*ELP1-TC6* cells (**Fig. 3F** and **Fig. S5**), FD patient-derived iPSC sympathetic neurons (**Fig. 3F** and **Fig. S6A-B**), and FD patient fibroblasts (**Fig. 3F** and **Fig. S7**).

Low-frequency off-target base editing events were detected at 9 sites in HEK293T-*ELP1-TC6* cells, 8 sites in FD iPSC-derived sympathetic neurons, and only a single site in FD patient fibroblasts (**Fig. 3G**). Importantly, all validated off-target editing events remained below 0.2% absolute editing frequency relative to paired untreated controls (**Fig. 3H**). Together, these findings demonstrate that TadCBEd-SpG paired with gRNA-C6 enables efficient correction of the *ELP1 T6C* mutation with minimal off-target editing across multiple human cell types.

To further investigate whether off-target activity could be mitigated through Cas9 enzyme engineering, we evaluated recently developed machine learning–guided Cas9 variants optimized for altered PAM selectivity via machine learning (*33*). We cloned and tested the TadCBEd version of SpCas9-LWRYEK, which has a preference for NGCG PAMs, and is therefore highly compatible with the gRNA-C6 target configuration.

Among the six off-target loci where we observed low but significant editing with TadCBEd-SpG, the LWRYEK variant reduced editing at two sites (OT2 and OT17) while largely preserving on-target editing activity (**Fig. 3I**). However, this variant also increased bystander editing at the intended *ELP1* target site (**Fig. 3I**). In addition, no substantial differences in off-target editing were observed at three loci (OT3, OT4, and OT20) relative to TadCBEd-SpG, whereas editing increased at one off-target site (OT19), which enconded NGCG PAM sequence compatible with LWRYEK’s PAM preference (**Fig. 3I**). Together, these findings suggest that customized PAM-selective Cas9 variants can modulate the specificity landscape of base editors, although improvements in certain off-target contexts may be accompanied by tradeoffs in bystander or PAM-compatible off-target editing. These results further support the continued development and usage of engineered Cas9 variants as a strategy to improve the therapeutic specificity of base editing.

### *In vivo* evaluation of ELP1 base editing and functional rescue in FD neuronal models

To evaluate the translational potential of our base editing strategy, we delivered intein-split TadCBEd-SpG paired with gRNA-C6 using two relevant AAV delivery approaches in the humanized *TgFD9* mouse model, which carries the human *ELP1* transgene harboring the FD-associated T6C mutation (*34*). The *TgFD9* mouse is perfectly suited to assess in vivo therapeutic modulation of human *ELP1* splicing, as it recapitulates the same tissue-specific mis-splicing observed in FD patients. However because these mice express normal endogenous murine *Elp1* gene, they do not exhibit neurological phenotypes (*34*).

We first tested localized ocular delivery using intravitreal administration of dual AAV2 vectors encoding the N- and C-terminal split base editor components in adult *TgFD9* mice, together with a GFP-expressing AAV reporter (**Fig. 4A**). Four to six weeks after injection, retinal cells were isolated and GFP-positive cells were enriched by fluorescence-activated cell sorting (FACS) to analyze cells that had likely received the co-administered vectors. Sequencing analysis of the sorted GFP-positive retinal cell population revealed efficient on-target editing, with some animals reaching up to ∼8% correction at the target cytosine and average editing levels of approximately 3% (**Fig. 4B**). Minimal bystander editing and indel formation were observed, consistent with the high precision of the SpG/gRNA-C6 configuration established *in vitro*.

**Fig. 4.**
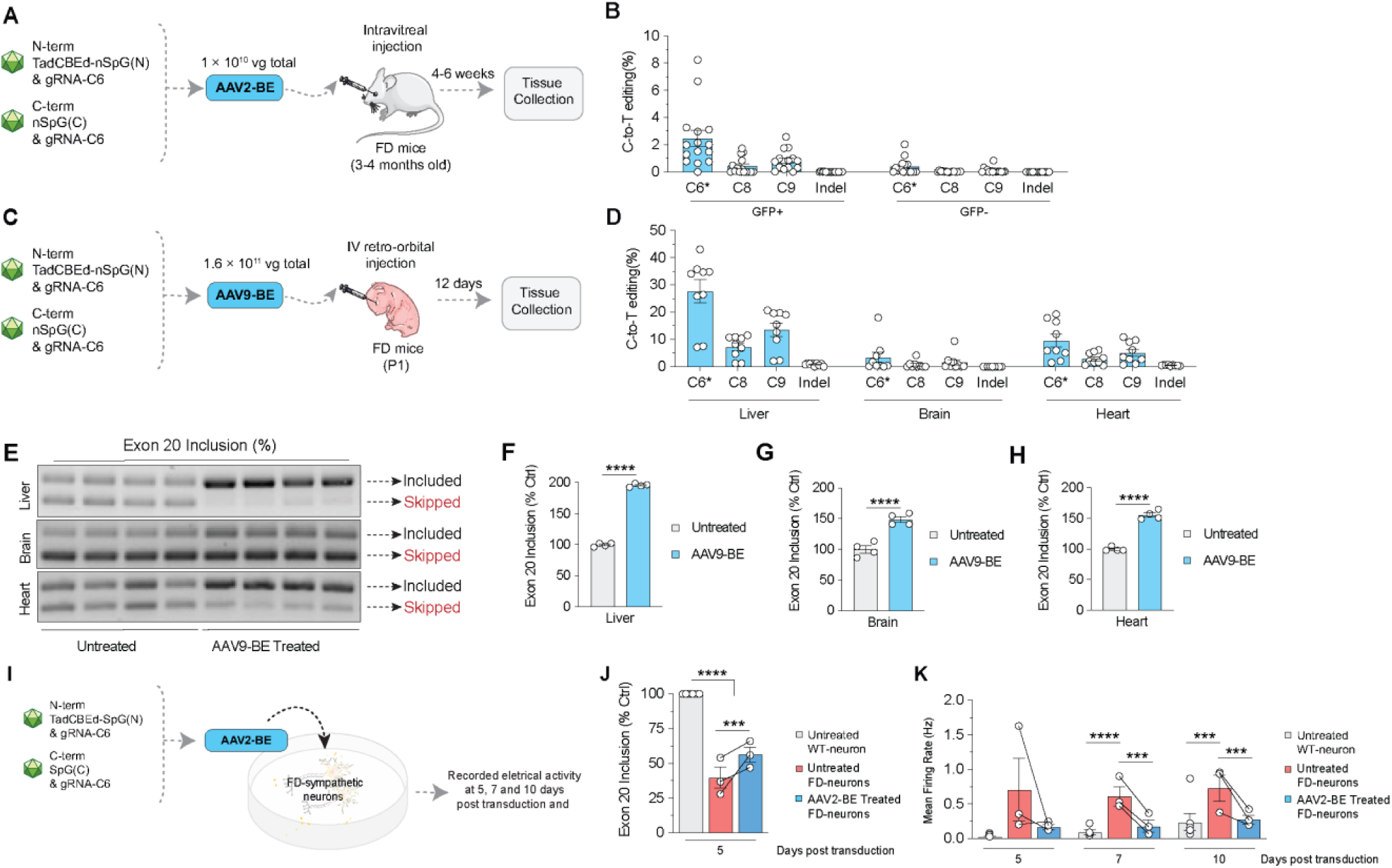
AAV-mediated delivery of intein-split base editors restores ELP1 splicing in vivo and rescues neuronal phenotypes in FD models. **(A)** Schematic of intravitreal delivery of dual AAV2 vectors encoding intein-split TadCBEd-SpG and gRNA-C6 into adult TgFD9 mice carrying the human ELP1 transgene harboring the FD-associated T6C mutation. Retinal cells were collected 4–6 weeks after injection and sorted based on GFP expression. **(B)** On-target and bystander C-to-T editing frequencies in GFP-positive and GFP-negative retinal cells following intravitreal AAV2-BE delivery. **(C)** Schematic of systemic retro-orbital intravenous administration of dual AAV9 vectors encoding intein-split TadCBEd-SpG and gRNA-C6 into neonatal (P1) TgFD9 mice. Tissues were collected 12 days after injection. **(D)** On-target and bystander editing frequencies in liver, brain, and heart tissues following systemic AAV9-BE delivery. Editing frequencies were quantified by targeted deep sequencing and analyzed using CRISPResso2. **(E)** Representative RT-PCR analysis of ELP1 exon 20 inclusion in liver, brain, and heart tissues from untreated and AAV9-BE-treated TgFD9 mice. Included and skipped ELP1 splice isoforms are indicated. (**F** to **H**) Quantification of exon 20 inclusion in liver (**F**), brain (**G**), and heart (**H**) tissues following systemic AAV9-BE treatment. **(I)** Schematic of AAV2-BE transduction of FD patient-derived iPSC sympathetic neurons and multielectrode array (MEA)-based neuronal activity recordings. **(J)** Quantification of ELP1 exon 20 inclusion in FD iPSC-derived sympathetic neurons following AAV2-BE treatment. **(K)** Mean firing rate measurements of healthy control neurons, untreated FD sympathetic neurons, and AAV2-BE-treated FD sympathetic neurons recorded at 5, 7, and 10 days after transduction. Data in panels **B**, **D**, and **F** to **K** represent mean ± SEM with individual data points shown; n = 3–6 independent biological replicates. Statistical analyses were performed as indicated in the Methods.

We next evaluated systemic delivery using retro-orbital intravenous administration of dual AAV9 vectors in neonatal (P1) *TgFD9* mice (**Fig. 4C**). Twelve days after injection, targeted sequencing demonstrated editing across multiple tissues, with the highest levels detected in liver (∼30% average editing with up to ∼45% observed), followed by brain (∼5% average editing) and heart tissues (∼10% average editing) (**Fig. 4D**). To determine whether these editing levels were sufficient to rescue *ELP1* splicing *in vivo*, we next analyzed *ELP1* exon 20 inclusion in tissues collected from AAV9-treated *TgFD9* mice. We observed a substantial increase in correctly spliced *ELP1* transcripts in liver, brain, and heart tissues compared to untreated *TgFD9* controls (**Fig. 4E-H**). Notably, despite relatively modest editing levels in the brain, *ELP1* exon 20 inclusion increased by approximately 50% following treatment (**Fig. 4G**), suggesting that partial correction of the mutant allele is sufficient to significantly increase WT ELP1 transcript level in the CNS.

To determine whether base editing could rescue molecular and functional phenotypes in FD patient-derived neurons, we differentiated FD iPSC into sympathetic neurons (*35, 36*). Sympathetic neurons were derived by direct differentiation of FD iPSCs through neural crest and sympathetic neuron progenitor stages (**Fig. S8A**). After 20 days of differentiation, mature FD iPSCs were differentiated into sympathetic neurons. Mature neurons were transduced with AAV2-BE vectors encoding the intein-split TadCBEd-SpG/gRNA-C6 system and analyzed 5 days later (**Fig. 4I**). Consistent with our *in vivo* findings, treated neurons exhibited increased *ELP1* exon 20 inclusion relative to untreated FD neurons (**Fig. 4J** and **Fig. S6C**).

To confirm that correction of *ELP1* splicing translated into functional improvement, we assessed neuronal activity using multielectrode array recordings at 5, 7, and 10 days following AAV transduction (**Fig. S8B**). As previously reported, untreated FD sympathetic neurons displayed marked spontaneous hyperactivity when compared to healthy control neurons (**Fig. 4K**). In contrast, AAV2-BE-treated FD neurons showed a substantial reduction in neuronal firing activity, approaching levels observed in control cultures across all examined time points (**Fig. 4K**). Together, these findings demonstrate that AAV-mediated base editing can restore *ELP1* splicing and rescue disease-relevant functional phenotypes in human FD sympathetic neurons.

## Discussion

Familial dysautonomia (FD) is a severe and progressive neurodegenerative disorder with no disease-modifying therapies, highlighting a critical unmet medical need. In this study, we developed a cytosine base editing strategy designed to precisely correct the FD pathogenic intronic splicing mutation. Through a systematic engineering campaign integrating SpCas9 and deaminase variants associated with guide RNA design, we identified base editor configurations that maximize on-target correction. Across multiple model systems, including human cells, patient-derived neurons, and a humanized mouse model, this approach restored *ELP1* exon 20 inclusion and improved disease-relevant phenotypes, establishing a foundation for a durable genome editing–based therapeutic strategy for FD.

A key conceptual advance of this work is the demonstration that base editors can be tailored to specific disease-causing sequences to achieve optimal precision and functional rescue. The *ELP1 T6C* mutation presents several inherent challenges, including limited PAM accessibility and the presence of adjacent cytosines that increase the risk of bystander editing. By leveraging PAM-relaxed Cas9 variants and systematically screening editing configurations, we identified distinct editor architectures that balance efficiency and precision. Notably, TadCBEd-SpG paired with gRNA-C6 achieved the highest editing efficiencies, whereas TadCBEd-SpRY paired with gRNA-C11 enabled precise editing with minimal bystander activity. Importantly, targeting splicing mutations in intronic regions offers a unique advantage because bystander edits would not affect protein coding. Although such edits have the potential to influence splicing and therefore must be carefully evaluated, in our study, bystander edits were in fact advantageous in enhancing *ELP1* exon 20 inclusion, further supporting the therapeutic potential of our approach. Thus, these findings support a broader paradigm in which genome editing enzymes are engineered for individual mutations, rather than relying on one-size-fits-all editing platforms (*21, 37*) .

Our results demonstrate that correction of the disease-causing mutation in a subset of cells is sufficient to restore neuronal function, revealing a favorable therapeutic threshold for FD. In patient-derived sympathetic neurons, ∼10% editing efficiency resulted in normalization of neuronal hyperactivity. This observation is consistent with our previous in vivo studies showing that even a modest increase in full-length *ELP1* expression is sufficient to rescue disease phenotype, including neuronal loss (*13, 38*). More broadly, these findings align with emerging evidence across genetic diseases that incomplete editing can yield substantial therapeutic benefit by restoring functional protein levels above a required threshold. (*21, 39*). In the present study, the disproportionate greater improvement in splicing correction relative to DNA editing efficiency suggests that even limited correction of the mutant allele can produce a substantially larger increase in correctly spliced *ELP1* transcripts. . From a translational perspective, this feature is highly advantageous, as it reduces the requirement for high editing efficiencies in tissues that are difficult to target, such as the central nervous system (CNS). However, future studies focused on improving delivery and alternative non-viral delivery platforms are highly desirable to further enhance precise editing.

A critical consideration for the clinical development of genome editing therapies is safety. We therefore performed a comprehensive assessment of off-target activity using GUIDE-seq2 and in silico prediction, followed by targeted validation across multiple human cell types. These analyses revealed minimal off-target editing, with all detected events occurring at very low frequencies. These findings are consistent with recent advances in base editor design that have improved specificity while maintaining high on-target activity (*40, 41*), and alternative use of novel PAM-specific enzymes, such as the LWRYEK variant demonstrated in this study (*33*).

In summary, we demonstrate that a customized base editing strategy can precisely and durably correct the genetic defect underlying FD, restore normal *ELP1* splicing, and rescue disease-relevant neuronal phenotypes. The finding that modest levels of editing level are sufficient for functional recovery provides a strong rationale for clinical development. More broadly, this work establishes a framework for the design of mutation-specific genome editing therapies and supports the application of base editing to a wide range of splicing disorders and neurological diseases driven by single-nucleotide variants.

## Methods

### Plasmids and oligonucleotides

gRNA target sequences, plasmids, and oligonucleotides used in this study are listed in **Tables S2-S4**. Newly generated plasmids will be deposited in Addgene. Base editor constructs were generated by cloning TadA-derived deaminase domains into pCMV-T7-ABEmax(7.10)-VRQR-P2A-EGFP (Addgene plasmid #140003) using isothermal assembly(42). Additional mutations were introduced using the Q5 Site-Directed Mutagenesis Kit (New England Biolabs). Human U6 promoter-driven gRNA expression plasmids were generated by cloning annealed oligonucleotides containing spacer sequences into BsmBI-digested pUC19-U6-BsmBI_cassette-SpCas9_gRNA (Addgene plasmid #65777). Intein-split base editor constructs were generated using Npu intein-based N- and C-terminal AAV backbones (Addgene plasmids #137177 and #137178). The N-terminal vector encoded the deaminase domain and gRNA expression cassette, whereas the C-terminal vector encoded the complementary Cas9 fragment and ELP1-targeting spacer sequences.

### HEK293T and fibroblast culture

HEK293T cells (ATCC) and FD patient fibroblasts (GM02343; Coriell Institute) were maintained in Dulbecco’s modified Eagle medium (DMEM) supplemented with 10% heat-inactivated fetal bovine serum (HI-FBS) and 1% penicillin-streptomycin. HEK293T cells were seeded at 2X10^4^ cells per well in 96-well plates and transfected approximately 20h later using TransIT-X2 (Mirus). Unless otherwise indicated, transfections contained 70 ng of base editor plasmid and 30 ng of guide RNA (gRNA) expression plasmid per well. For plasmid titration experiments, an inert stuffer plasmid was added to maintain a constant total DNA amount. Genomic DNA was collected 72 h after transfection using a proteinase K-based lysis protocol and stored at −20°C until analysis. Clonal HEK293T-ELP1-TC6 cell lines were generated by limiting dilution and expansion of single-cell-derived colonies. Cell cultures were routinely screened for mycoplasma contamination using PCR based kits and tested negative for mycoplasma.

### Human pluripotent stem cell (hPSC) lines

hPSC experiments were performed using two female cell lines: H9 human embryonic stem cells (WA09; WiCell; NIH registry #0062) and S2 induced pluripotent stem cells (iPSCs) derived from a patient with familial dysautonomia (FD). The S2 iPSC line was previously generated and characterized from FD patient fibroblasts (GM04899; Coriell Institute) (*42*). Detailed hPSC maintenance and characterization procedures have been described previously (*42, 43*). hPSCs were maintained in Essential 8 medium (Gibco) on vitronectin-coated culture plates (Thermo Fisher Scientific) and passaged using EDTA according to the manufacturer’s instructions.

### In vitro differentiation of hPSCs into sympathetic neurons

hPSCs were differentiated into sympathetic neurons as previously described (*44*). Briefly, hPSCs were induced toward a neural crest fate using sequential modulation of BMP, TGF-β, and WNT signaling pathways. Neural crest cells were subsequently aggregated into three-dimensional spheroid cultures and differentiated into sympathetic neuron progenitors. On day 14, progenitors were dissociated and replated onto polyornithine-, laminin-, and fibronectin-coated culture plates for terminal differentiation and maturation. Neurons were maintained in Neurobasal medium supplemented with N2, B27, L-glutamine, GDNF, BDNF, NGF, ascorbic acid, dbcAMP, and retinoic acid as previously described (*44*). Independent biological replicates were defined as differentiations initiated from separate hPSC cultures at least 3 days apart or from independently thawed hPSC stocks.

### AAV2-BE transduction in iPSC-derived sympathetic neurons and neuronal activity analysis

FD iPSC-derived sympathetic neurons were transduced with AAV2 vectors encoding the intein-split TadCBEd-SpG base editor and gRNA-C6 on day 21 of differentiation. Viral vectors were applied directly to neuronal cultures for 6 h, after which fresh differentiation medium was added. Twenty-four hours later, virus-containing medium was replaced with fresh medium, and neurons were maintained under standard differentiation conditions.

Neuronal morphology and viability were monitored by bright-field microscopy following transduction. GFP fluorescence was assessed using a Lionheart FX automated microscope (BioTek) to confirm transduction efficiency. For genomic DNA analysis, neurons were harvested 5 days after transduction, washed with PBS, collected by gentle scraping, and stored at −80°C until processing.

Neuronal activity was evaluated as previously described (*44*). Briefly, sympathetic neuron progenitors were plated on polyornithine-, laminin-, and fibronectin-coated 96-well multielectrode array (MEA) plates (Axion Biosystems) on day 14 of differentiation and transduced as described above. Spontaneous neuronal activity was recorded at the indicated time points using an Axion Maestro Pro system and analyzed using the neural detection mode.

### Next-generation sequencing (NGS) and data analysis

Genome editing efficiencies were quantified by targeted next-generation sequencing (NGS) using a two-step PCR-based Illumina library preparation workflow adapted from previous studies (*27*). Briefly, genomic regions of interest were amplified from approximately 50 ng of genomic DNA using Q5 High-Fidelity DNA Polymerase (New England Biolabs) and locus-specific primers (**Table S4**). Amplicons were purified using paramagnetic beads (*45, 46*), followed by a second PCR to incorporate sample-specific barcodes and Illumina adapter sequences. Libraries were purified, quantified by capillary electrophoresis (QIAxcel, Qiagen) and qPCR (KAPA Library Quantification Kit, Roche), normalized, pooled, and sequenced on an Illumina MiSeq platform using 300-cycle v2 chemistry.

Sequencing data were analyzed using CRISPResso2 (*47*). Nuclease-mediated editing was quantified using standard CRISPResso2 alignment and indel analysis parameters: CRISPResso -r1 READ1 -r2 READ2 --amplicon_seq--amplicon_name --guide_seq GUIDE -w 20 --cleavage_offset -10. Base editing outcomes were quantified using the CRISPResso2 base editor workflow configured to detect C-to-T conversions (or A-to-G) conversations: CRISPResso -r1 READ1 -r2 READ2 --amplicon_seq --guide_seq GUIDE -w 20 --cleavage_offset -10 --base_editor_output --conversion_nuc_from C --conversion_nuc_to T --min_frequency_alleles_around_cut_to_plot 0.001. Editing frequencies were calculated from aligned sequencing reads and reported as the percentage of modified alleles at the target nucleotide unless otherwise indicated.

### Off-target analysis via GUIDE-seq2

Genome-wide off-target activity was assessed using GUIDE-seq2, an updated version of the GUIDE-seq method (*30–32*). HEK293T-ELP1-T6C cells were transfected with nuclease expression plasmids, guide RNA (gRNA) plasmids, and a double-stranded oligodeoxynucleotide (dsODN) integration tag. Genomic DNA was isolated 72 h after transfection using the DNAdvance Kit (Beckman Coulter). Integration of the GUIDE-seq dsODN at the on-target locus was quantified by targeted next-generation sequencing and analyzed using CRISPResso2 (*47*). The frequency of dsODN integration was calculated as the proportion of sequencing reads containing the integrated tag relative to total aligned reads at the target locus.

Genome-wide GUIDE-seq2 libraries were generated essentially as described previously, with modifications based on the GUIDE-seq2 workflow (*30–32*). Briefly, genomic DNA was subjected to Tn5-mediated tagmentation using adapters containing sample barcodes and unique molecular identifiers (UMIs) (**Table S4**). Libraries enriched for dsODN integration sites were generated by PCR amplification using strand-specific primers, size selected, quantified, and sequenced on an Illumina NextSeq 1000/2000 platform.

Sequencing reads were demultiplexed and downsampled to equal read depths across samples for comparisons performed with the same gRNA. Candidate off-target sites were identified using the GUIDE-seq2 analysis pipeline (https://github.com/tsailabSJ/guideseq/tree/V2) with the maximum mismatch parameter set to six. The most prominent candidate off-target sites identified by GUIDE-seq2 and Cas-OFFinder were subsequently validated by targeted next-generation sequencing using locus-specific primers listed in **Table S5**.

### AAV production

Plasmids encoding intein-split TadCBEd-SpG and gRNA-C6 for packaging into AAV2 and AAV9 vectors were generated as previously described (*20, 48, 49*) and are listed in **Table S3**. Recombinant AAV vectors were produced by UMass Chan Medical School Viral Vector Core. Vector genome titers were determined by droplet digital PCR (ddPCR), and vector purity was assessed by SDS-PAGE.

### Mice and AAV-BE treatment *in vivo*

The humanized TgFD9 mouse model carrying the human *ELP1* transgene harboring the FD-associated intron 20 T6C mutation was previously described (*34*). TgFD9 mice and wild-type littermate controls were maintained on a mixed C57BL/6J and C57BL/6N background. Both male and female animals were included in all studies. Mice were housed under standard conditions with ad libitum access to food and water on a 12h light/dark cycle. All animal experiments were approved by the Massachusetts General Hospital Institutional Animal Care and Use Committee and performed in accordance with NIH guidelines.

Genotyping was performed using genomic DNA isolated from tail biopsies and PCR-based detection of endogenous *Elp1* alleles and the TgFD9 transgene using primers listed in **Table S4**.

For retinal studies, adult TgFD9 mice received intravitreal injections of dual AAV2 vectors encoding intein-split TadCBEd-SpG and gRNA-C6 under isoflurane anesthesia. A total volume of 1 μL containing the indicated vector combination (45% N-terminal vector, 45% C-terminal vector, and 10% AAV2-eGFP control vector) was administered at doses of 1.5 × 10^9 or 1 × 10^10 vector genomes (vg) per eye. Animals received topical antibiotic treatment and postoperative analgesia following injection.

For systemic delivery studies, neonatal (P1) TgFD9 mice received dual AAV9 vectors encoding intein-split TadCBEd-SpG and gRNA-C6 by temporal facial vein injection as previously described (*50*). N-terminal and C-terminal vectors were administered at total doses of 3 × 10^10^ diluted in phosphate-buffered saline (PBS). Pups were monitored throughout the procedure and returned to the dam following recovery.

### Isolation of retinal cell suspension

Retinas were dissected and dissociated into single-cell suspensions using a papain-based enzymatic digestion protocol as previously described (*51*). Retinal tissue was incubated in papain solution containing DNase I, followed by quenching with ovomucoid-containing buffer and mechanical trituration. Dissociated cells were filtered through a 20-μm cell strainer to remove residual tissue fragments and centrifuged at 300 × g for 10 min at 4°C. The resulting single-cell suspensions were used for downstream fluorescence-activated cell sorting and genomic DNA analysis.

### Extraction of gDNA and RNA from mouse tissues

Genomic DNA was isolated from frozen mouse tissues using the Agencourt DNAdvance DNA purification system (Beckman Coulter) according to the manufacturer’s instructions. DNA concentrations were determined using a NanoDrop spectrophotometer (Thermo Fisher Scientific).

For RNA analyses, tissues were rapidly dissected, snap-frozen in liquid nitrogen, and homogenized in TRI Reagent (Molecular Research Center) using a TissueLyser (Qiagen). Total RNA was extracted according to the manufacturer’s protocol, and RNA concentration and purity were assessed by NanoDrop spectrophotometry. cDNA synthesis was performed from 1 μg of total RNA using random primers (Promega) and SuperScript III reverse transcriptase (Invitrogen) as previously described (*13, 14*).

### RT-PCR analysis of *ELP1* splicing

Alternative splicing of human *ELP1* transcripts was assessed by RT-PCR using cDNA generated from 100 ng of total RNA. Amplification was performed using GoTaq Green Master Mix (Promega) and human-specific *ELP1* primers (**Table S4**). PCR products were resolved on 1.5% agarose gels and visualized by ethidium bromide staining. The relative abundance of full-length and exon 20-skipped (*Δ20*) *ELP1* transcripts was quantified using ImageJ based on integrated band intensities, as previously described (*13, 14, 34, 52*). Exon 20 inclusion was calculated as the percentage of full-length *ELP1* transcript relative to the total *ELP1* transcript signal.

### RT-qPCR analysis of *ELP1* transcripts

Expression of full-length (FL) and exon 20-skipped (Δ20) *ELP1* transcripts was quantified by TaqMan-based RT-qPCR using the CFX384 Touch Real-Time PCR Detection System (Bio-Rad) and One-Step RT-qPCR reagents (Bio-Rad). Primer and probe sequences are provided in **Table S4**. RT-qPCR reactions were performed using RNA equivalent to 25 ng of input material per reaction. Transcript abundance of *FL ELP1*, *Δ20 ELP1*, and *GAPDH* was quantified using isoform-specific TaqMan assays. Expression values were normalized to *GAPDH* and vehicle-treated controls and are presented as fold change relative to control samples. Data were analyzed using SDS software.

### Statistical analysis

Data are presented as mean ± SEM unless otherwise indicated. Statistical analyses were performed using GraphPad Prism 10 (GraphPad Software) for mouse studies and validation of off-target validation. Comparisons between two groups were performed using unpaired two-tailed Student’s t tests. Comparisons involving three groups were analyzed using one-way analysis of variance (ANOVA) followed by unpaired or paired two-tailed Student’s t tests for multiple-comparison correction tests. The number of independent biological replicates (n) is provided in the figure legends. Differences were considered statistically significant when P < 0.05.

### Data availability

Primary datasets will be available in **Table S6**. Sequencing datasets will be deposited with the NCBI Sequence Read Archive (SRA). Plasmids from this study will be made available through Addgene.

## Supporting information

Table S6 - Primary datasets

Table S5 - Off target validation analysis

Table S3 - Plasmids

Table S2 - gRNA target sites

Table S4 - Oligonucleotides

Table S1 - GUIDE-seq2 and Cas-OFFinder results

FD_BE_SupMaterials

## Acknowledgements

We acknowledge the following individuals for their technical contributions: Sabyasachi Das (in vivo injections and tissue collection) and Sydnee Dymock (mouse colony maintenance). We thank Dr. Alejandra Gonzalez-Duarte and Dr. Horacio Kaufmann of the Dysautonomia Treatment and Evaluation Center at New York University Medical School for their helpful discussions. This work was funded by National Institutes of Health (NIH) R21EY037018 (to C.R.R.A and E.M.) and R01NS142225 (to E.M.) and by Familial Dysautonomia Foundation Clare and Philip Wexler Early-Stage Career Investigator Award (to C.R.R.A and E.M.). C.R.R.A was also supported by a Charles A. King Trust Postdoctoral Research Fellowship, Bank of America, N.A., Co-Trustees, a James L. and Elisabeth C. Gamble Endowed Fund for Neuroscience Research / Mass General Neuroscience Transformative Scholar Award, a MGH Physician/Scientist Development Award, and a NIH grant K01NS134784. B.P.K. was supported by an MGH Howard M. Goodman Fellowship, the Kayden-Lambert MGH Research Scholar Award 2023-2028, and NIH grants DP2CA281401, P01HL142494 and R01NS125353. A.C. is supported by the Knights Templar Eye Foundation career starter grant and funding from the Familial Dysautonomia Foundation. E.M. was supported by the HMS Ruth and Maurice Freeman Award for Pain Research and NIH grants R01NS124561 and R01NS095640. S.A.S. was supported by the NIH grant R01NS095640. This work was supported by NIH (1R01NS114567-01A1) to N.Z., supporting J.A.

## Author contributions

S.Y., A.C., B.P.K., S.A.S, E.M. and C.R.R.A. conceived of and designed the study. All authors designed, performed, or supervised experiments, and/or analyzed data. S.Y. and D.R.R performed plasmid cloning, cell culture experiments with HEK293T cells, NGS assays, and additional molecular experiments. K.K., S. P. and P. P performed splicing analysis. J.A. and N.Z. designed, conducted and analyzed experiments using iPSCs derived FD neurons. A.C., K.K., and S.P. conducted the *in vivo* experiments using *TgFD9* mice. S.Y., E.M. and C.R.R.A wrote the manuscript with contributions and revisions from all authors.

## Competing interests

B.P.K., E.M. and C.R.R.A are inventors on a patent application filed by Mass General Brigham (MGB) that describes genome editing technologies to treat FD. B.P.K. and C.R.R.A. are inventors on additional patents or patent applications filed by MGB that describe genome engineering technologies. S.A.S. is an inventor on several U.S. and foreign patents and patent applications assigned to the Massachusetts General Hospital, including U.S Patents 8,729,025 and 9,265,766, both entitled “Methods for altering mRNA splicing and treating familial dysautonomia by administering benzyladenine,” filed on August 31, 2012 and May 19, 2014 and related to use of kinetin; and U.S. Patent 10,675,475 entitled, “Compounds for improving mRNA splicing” filed on July 14, 2017 and related to use of BPN-15477. E.M. and S.A.S. are inventors on an International Patent Application Number PCT/US2021/012103, assigned to Massachusetts General Hospital and entitled “RNA Splicing Modulation” related to use of BPN-15477 in modulating splicing. B.P.K. is or was a consultant for Jumble Therapeutics, Foresite Labs, Generation Bio, Novartis Venture Fund, and Foresite Capital, and is on the scientific advisory boards of Life Edit Therapeutics and Prime Medicine. B.P.K. has a financial interest in Prime Medicine, Inc., a company developing therapeutic CRISPR-Cas technologies for gene editing. E.M. is a consultant for ReviR Therapeutics. C.R.R.A was a consultant for Ilios Therapeutics and Biogen and holds stocks in publicly traded companies developing gene therapies. B.P.K. and C.R.R.A interests were reviewed and are managed by MGH and MGB in accordance with their conflict-of-interest policies. C.R.R.A interests were also reviewed and are managed by the University of Pittsburgh in accordance with their conflict-of-interest policies. N.Z. is the founder and owner of Neela Cell Therapeutics, LLV. All other authors declare no competing interests.

## Notes

### Summary of Updates

In this version, we have revised the text throughout to improve clarity and readability. Figures also have been updated to improve labelling.

