## Supplementary material for "Engineered CRISPR-Base Editors as a Permanent Treatment for Familial Dysautonomia": FD_BE_SupMaterials

### Supplementary Materials


<u>Supplementary Tables</u> *Attached separately*

**Table S1**: GUIDE-seq2 and Cas-OFFinder results

**Table S2**: gRNA target sites

**Table S3**: Plasmids

**Table S4**: Oligonucleotides and probes

**Table S5**: Off-target validation analysis

**Table S6**: Primary datasets


***NOTE:*** All Supplementary Tables are attached separately as .xlsx files.


<u>Supplementary Figures and Legends</u>

**Figures S1-S8** *pages 2-10*

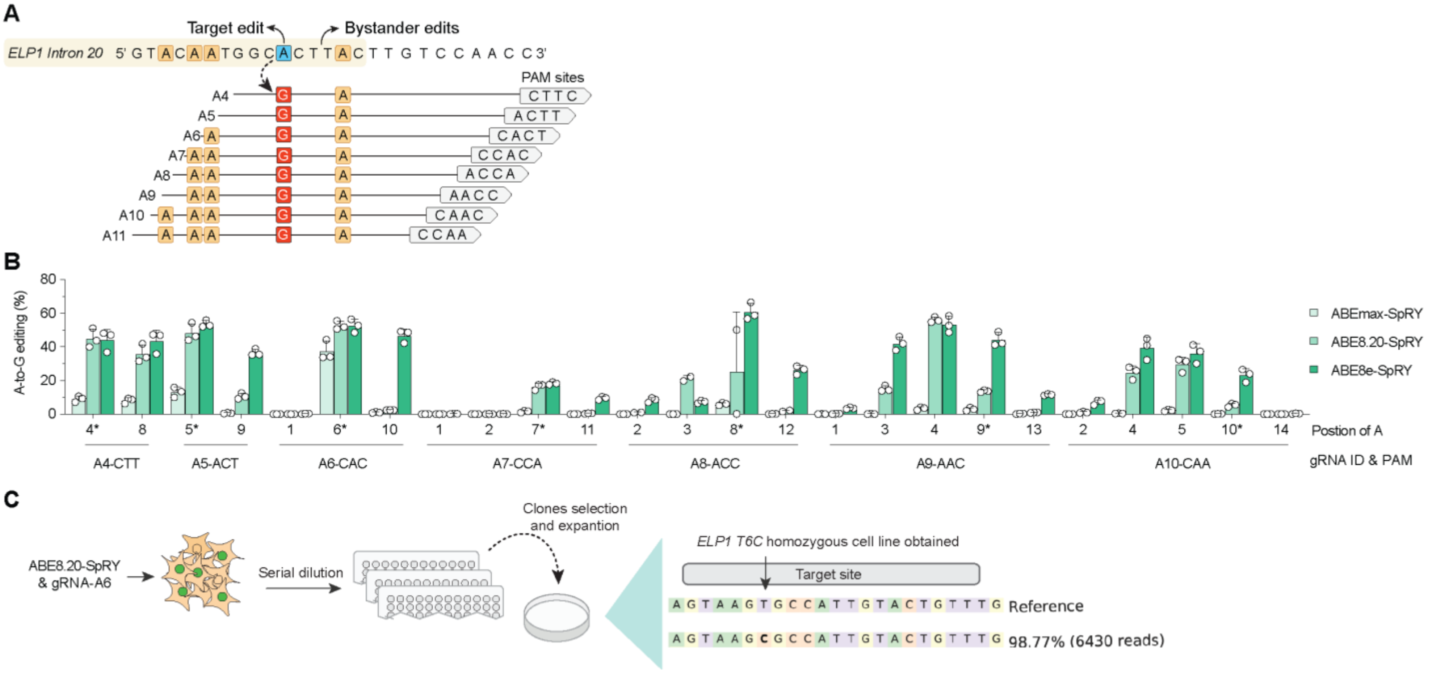


**Fig. S1. Generation of a HEK293T cell line harboring the ELP1 T6C mutation**

**(A** and **B)** Introduction of the ELP1 T6C mutation in HEK293T cells using adenine base editors (ABEs) fused to nSpRY and paired with multiple guide RNAs (gRNAs). Editing frequencies were quantified by targeted deep sequencing and analyzed using CRISPResso2.

**(B)** Experimental workflow used to generate clonal HEK293T-ELP1-TC6 cell lines. Cells were transfected with ABE8.20-nSpRY and gRNA-A6, followed by single-cell isolation, clonal expansion, and genotyping by targeted sequencing.

Data represent mean ± SEM with individual data points shown; $n$ = 3 independent biological replicates.

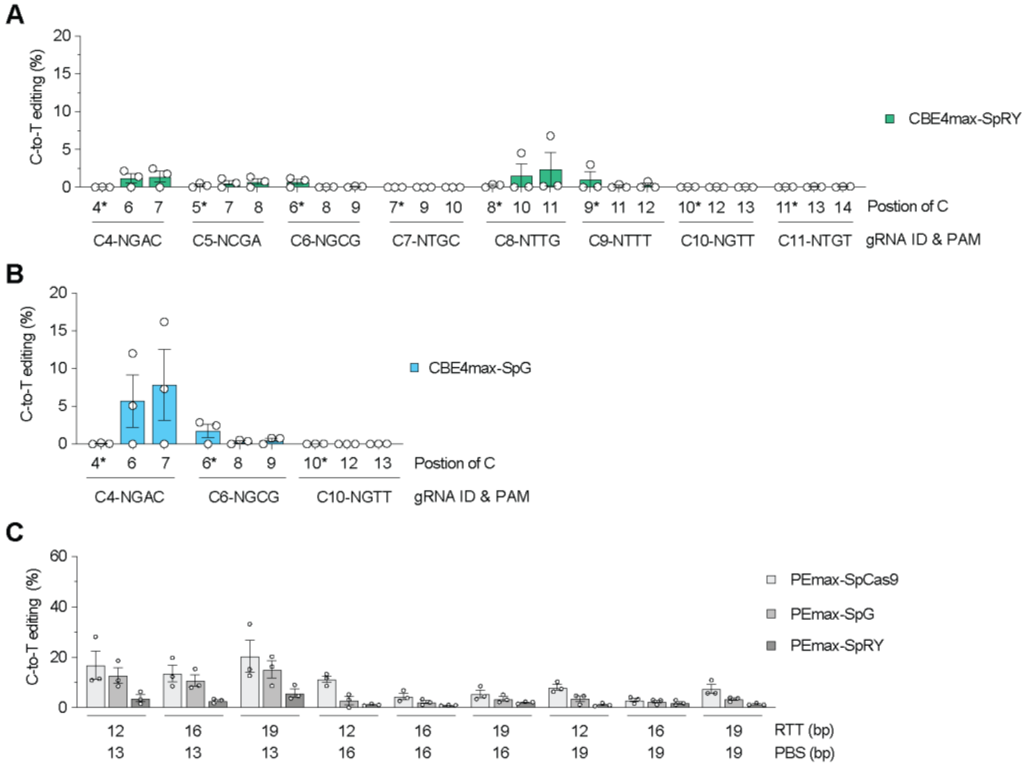


**Fig. S2. Correction of the ELP1 T6C mutation using BE4max and prime editing platforms.**

**(A** and **B)** C-to-T editing efficiencies in HEK293T-ELP1-TC6 cells using BE4max fused to nSpG (**A**) or nSpRY (**B**) paired with gRNAs targeting the ELP1 locus. Editing frequencies were determined by targeted deep sequencing and analyzed using CRISPResso2.

**(C)** Prime editing of the ELP1 T6C mutation using PEmax fused to WT SpCas9, nSpG, or nSpRY. Multiple reverse-transcription template (RTT) and primer-binding site (PBS) configurations were evaluated together with a secondary nicking guide RNA targeting the non-edited strand.

Data represent mean ± SEM with individual data points shown; *n* = 3 independent biological replicates.

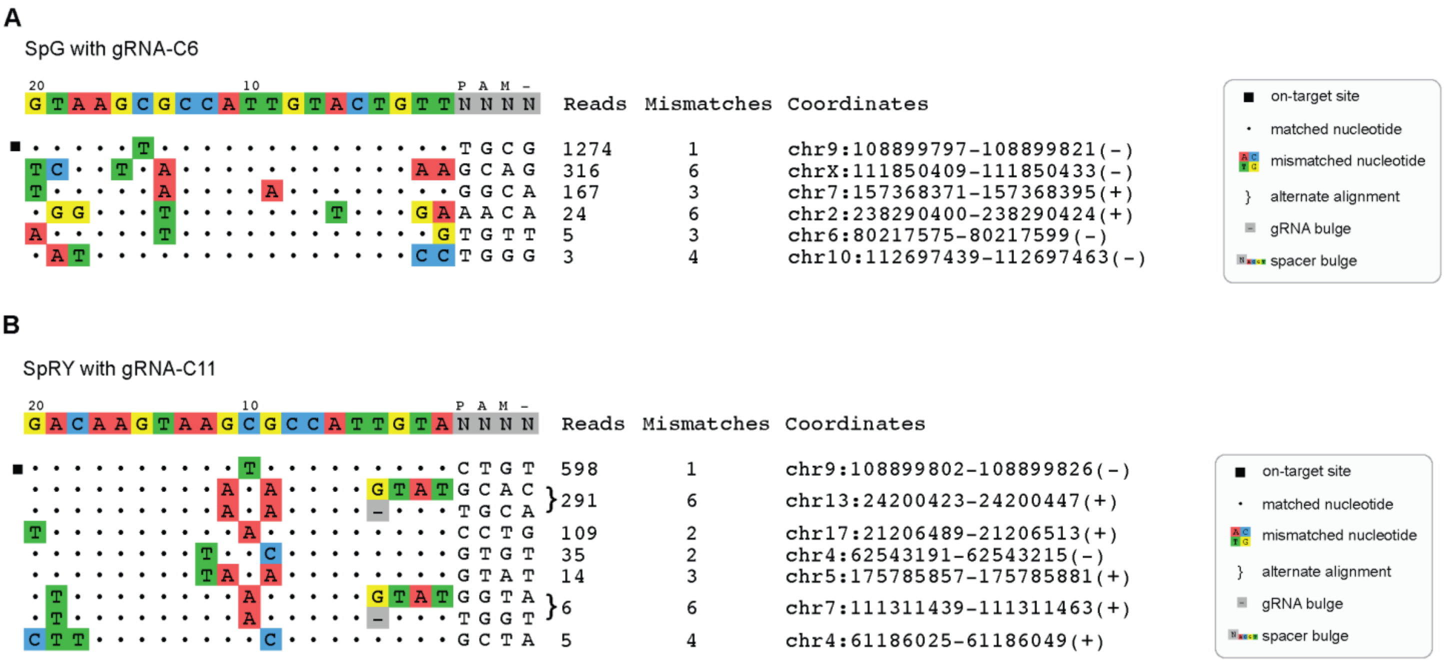


**Fig. S3. Identification of candidate off-target sites using GUIDE-seq2.**

**(A and B)** Rank-ordered GUIDE-seq2 profiles obtained from HEK293T-ELP1-TC6 cells transfected with SpG nuclease and gRNA-C6 (**A**) or SpRY nuclease and gRNA-C11 (**B**). Mismatched nucleotides relative to the on-target spacer sequence are highlighted. GUIDE-seq2 read counts corresponding to consolidated unique molecular events are shown adjacent to each site.

Data represent pooled sequencing libraries generated from *n* = 3 independent biological replicates.

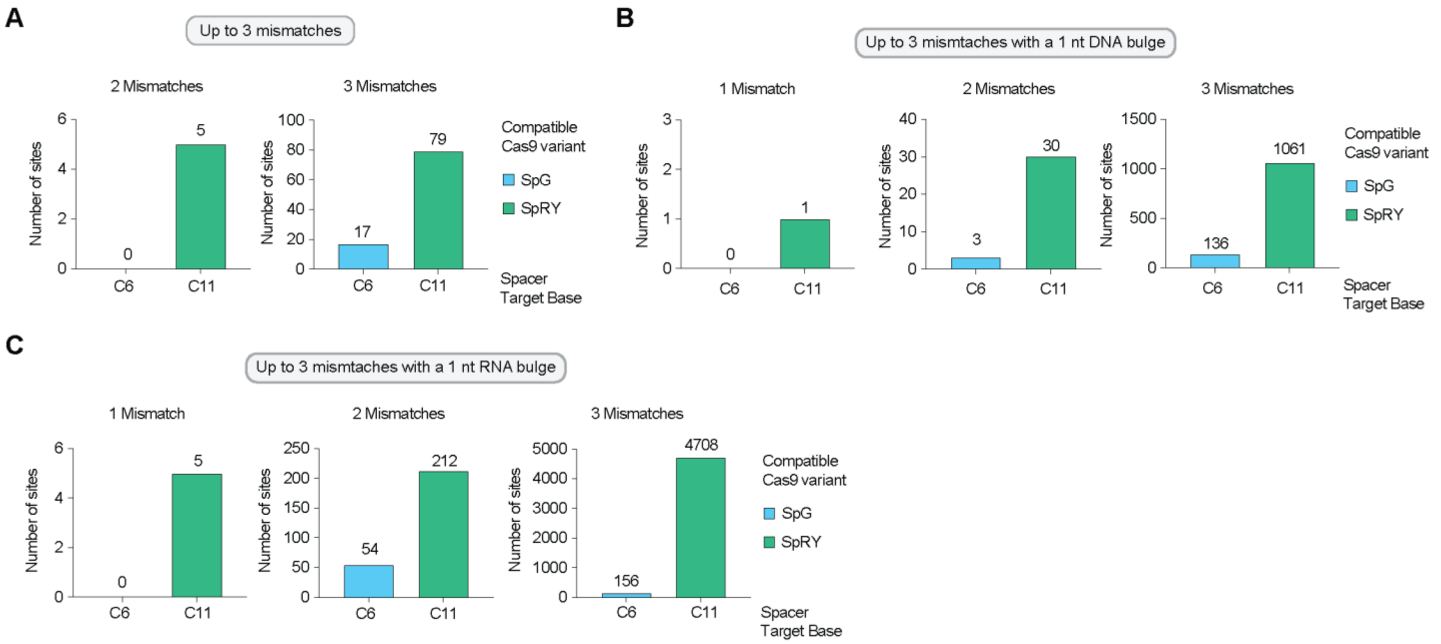


**Fig. S4. Computational prediction of potential off-target sites.**

**(A–C)** Cas-OFFinder prediction of candidate off-target sites for gRNA-C6 and gRNA-C11. Sites were identified allowing up to three mismatches (**A**), three mismatches plus a 1-nt DNA bulge (**B**), or three mismatches plus a 1-nt RNA bulge (**C**).

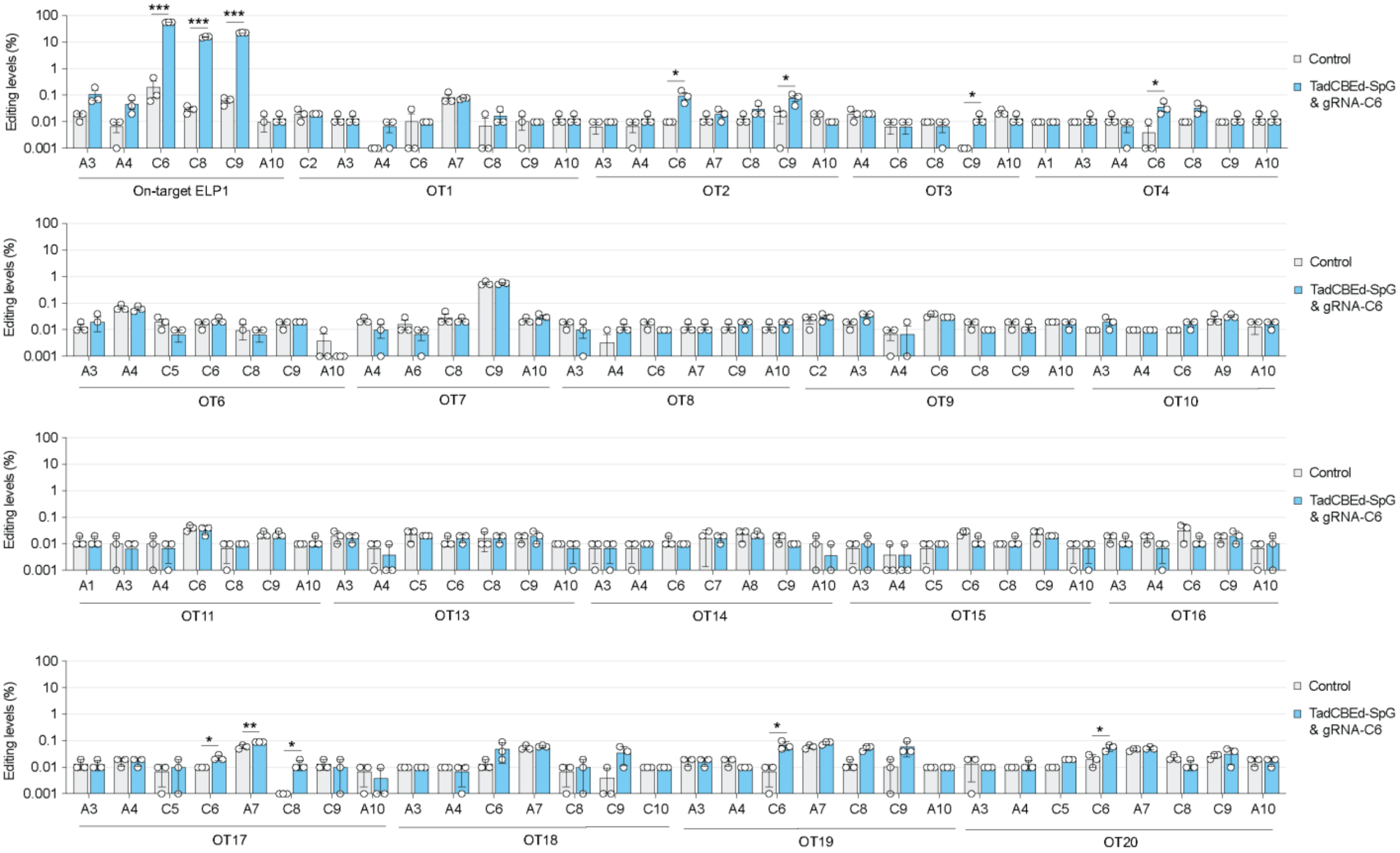


**Fig. S5. Validation of on-target and off-target editing in HEK293T-ELP1-TC6 cells.**

Targeted deep sequencing analysis of the ELP1 on-target locus and 20 nominated off-target sites in untreated HEK293T-ELP1-TC6 cells or cells treated with TadCBEd-nSpG and gRNA-C6. Editing frequencies for all cytosines and adenines within positions 1–10 of the protospacer sequence were quantified using CRISPResso2. Off-target sites were nominated by GUIDE-seq2 and Cas-OFFinder analyses. Genomic regions corresponding to OT5 and OT12 could not be amplified or sequenced.

Data represent n = 3 independent biological replicates.

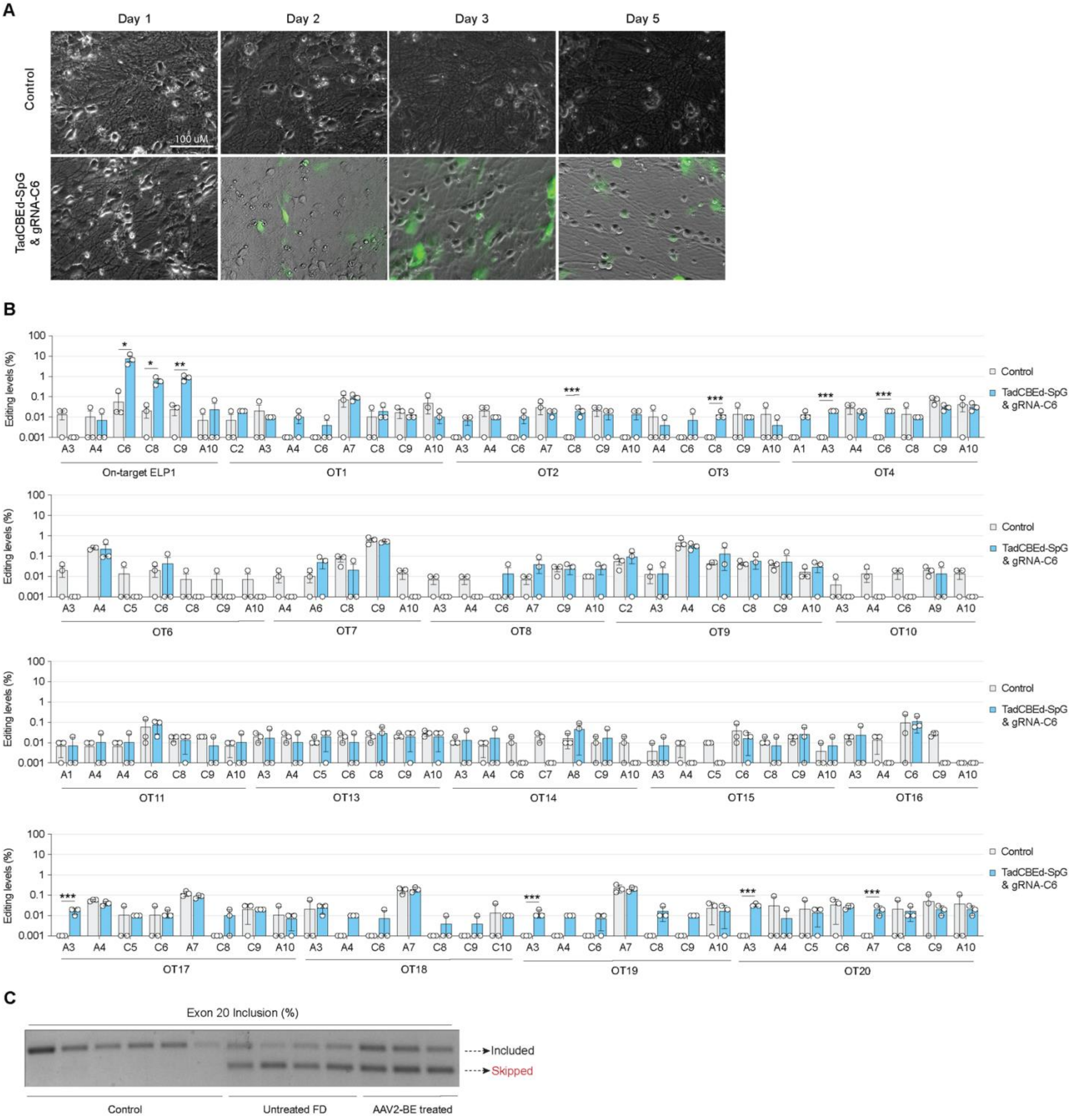


**Fig. S6. Validation of on-target and off-target editing in FD iPSC-derived sympathetic neurons.**

**(A)** Representative bright-field image of untreated FD iPSC-derived sympathetic neurons and GFP fluorescence image of neurons transduced with AAV2-BE.

**(B)** Targeted deep sequencing analysis of the ELP1 on-target locus and nominated off-target sites in untreated neurons or neurons transduced with AAV2-BE expressing TadCBEd-nSpG and gRNA-C6. Editing frequencies for all cytosines and adenines within positions 1–10 of the protospacer sequence were quantified using CRISPResso2. OT5 and OT12 failed amplification or sequencing.

**(C)** Representative RT-PCR analysis of ELP1 exon 20 inclusion in untreated and AAV2-BE-transduced FD iPSC-derived sympathetic neurons.

Data represent *n* = 3 independent biological replicates.

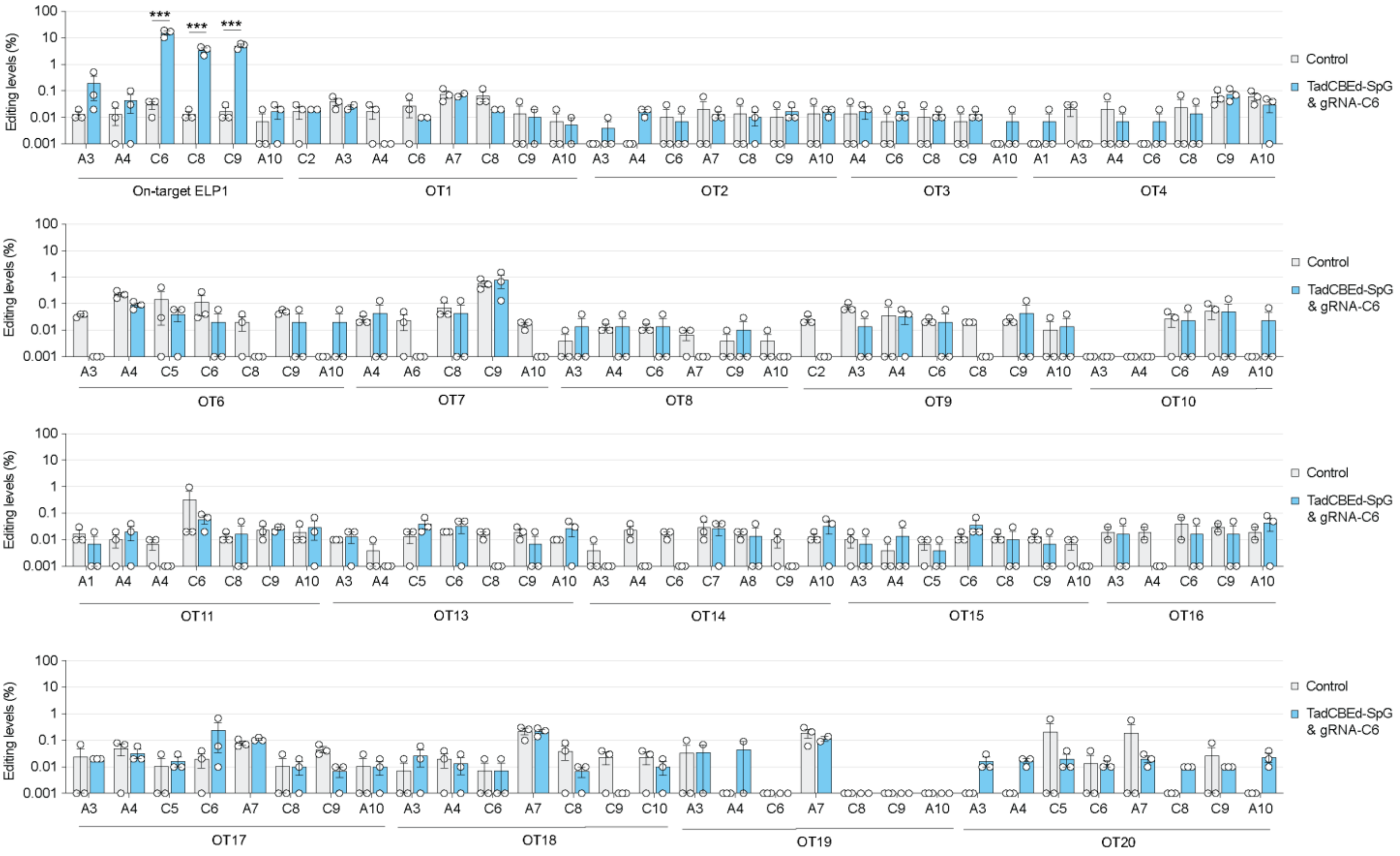


**Fig. S7. Validation of on-target and off-target editing in FD patient fibroblasts.**

Targeted deep sequencing analysis of the ELP1 on-target locus and nominated off-target sites in untreated FD patient fibroblasts or fibroblasts treated with TadCBEd-nSpG and gRNA-C6. Editing frequencies for all cytosines and adenines within positions 1–10 of the protospacer sequence were quantified using CRISPResso2. Candidate off-target sites were selected from GUIDE-seq2 and Cas-OFFinder analyses. OT5 and OT12 failed amplification or sequencing.

Data represent *n* = 3 independent biological replicates.

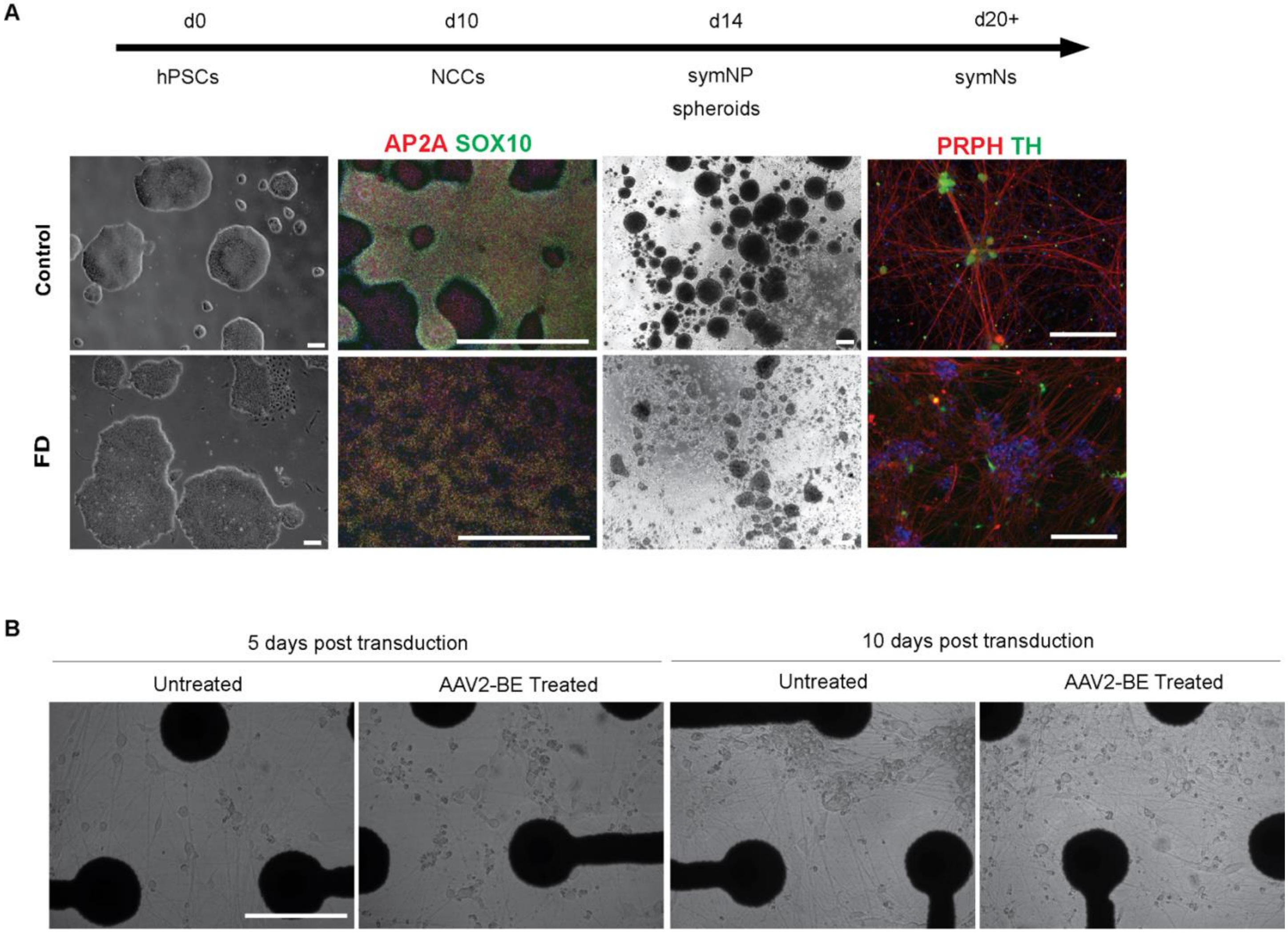


**Fig. S8. Sympathetic neurons derived by direct differentiation of FD iPSCs through neural crest and sympathetic neuron progenitor stages.**

**(A)** Healthy control and FD hPSCs were directly differentiated into neural crest cells (NCCs), positive for transcription factor AP2A and SOX10. FD iPSCs have impaired differentiation efficiency at d10 NCC stage, but a sympathetic neuron progenitor (symNP) spheroid phase allows for expansion and purification of the NCC population. After 20 days of differentiation, mature sympathetic neurons (symNs) express peripherin (PRPH) and tyrosine hydroxylase (TH), confirming symN identity. Scale bar represents 200 μm.

**(B)** Morphology of FD iPSC-derived sympathetic neurons following AAV2-BE transduction. Representative bright-field images of untreated and AAV2-BE-transduced FD iPSC-derived sympathetic neurons cultured on multielectrode array (MEA) plates at 5 and 10 days after transduction. Scale bars, 200 μm.
